# Incremental improvements in tractometry-based brain-age modeling with deep learning

**DOI:** 10.1101/2023.03.02.530885

**Authors:** Ariel Rokem, Joanna Qiao, Jason D. Yeatman, Adam Richie-Halford

## Abstract

Multivariate measurements of human brain white matter (WM) with diffusion MRI (dMRI) provide information about the role of WM in a variety of cognitive functions and in brain health. Statistical models take advantage of the regularities in these data to make inferences about individual differences. For example, dMRI data provide the basis for accurate brain-age models – models that predict the chronological age of participants from WM tissue properties. Deep learning (DL) models are powerful machine learning models, which have been shown to provide benefits in many multivariate analysis settings. We investigated whether DL would provide substantial improvements for brain-age models based on dMRI measurements of WM in a large sample of children and adolescents. We found that some DL models fit the data better than a linear baseline, but the differences are small. In particular, recurrent neural network architectures provide up to ∼6% improvement in accuracy. This suggests that information about WM development is mostly accessible with linear models, and does not require the additional invariance and non-linearity offered by DL models. However, in some applications this incremental improvement may prove critical. We provide open-source software that fits DL models to dMRI data (https://yeatmanlab.github.io/AFQ-Insight).

## Introduction

Neuroimaging data is very high-dimensional. In many neuroimaging applications, we reduce the dimensionality of the data using our knowledge of the anatomical structure of the brain, and simplify it by summarizing a lower-dimensional representation in a “tidy” tabular format that is amenable to standard statistical analysis approaches ^1^. In the field of machine learning, this is known as *feature engineering*. One example of anatomically-informed feature engineering in neuroscience is *tractometry* ^*2*^, which focuses on quantification of the tissue properties of brain white matter pathways. These pathways form the backbone of brain networks, and their physical properties are important for brain health and cognitive functions ^3^. Tractometry uses diffusion MRI (dMRI) measurements, which are sensitive to the random motion of water molecules within brain tissue in a location- and direction-specific manner. In densely packed white matter, water diffusion is less restricted along the length of bundles of axons that travel coherently through the white matter than across their boundaries. More generally, the diffusion process probes the cellular structure in a voxel. This means that the patterns of diffusion within each voxel can serve as cues for computational tractography algorithms that trace the trajectory of axonal bundles through the white matter. Anatomical knowledge can be applied to group these trajectories into estimates of the trajectories of the major white matter tracts. In addition, the patterns of diffusion can be summarized in every voxel in the brain to characterize physical properties of the tissue. These tissue properties are summarized as a scalar quantity in each voxel – such as the mean diffusivity, MD, or the fractional anisotropy of diffusion, FA – and can then be associated with different positions along the length of the tracts that were derived in tractography. These measurements are then organized into a “tidy” tabular format, where each column corresponds to a specific location along the length of each pathway or tract, and the tissue property that was measured in that location, and each row are the measurements from one individual ^4,5^. While the data can be organized into a table in this way, it does retain some of its original structure, because neighboring columns in a table into which this data is organized are also neighboring locations in the brain. In previous work, we capitalized on this knowledge to fit models that relate white matter tissue properties to phenotypic variance among the subjects. We used the grouping of the features into anatomical units to fit group-regularized models ^5^. However, while this previous approach distinguished between features that belong to the same anatomical group and features that belong to distinct anatomical groups, it did not differentiate between features that are in closer or farther proximity within each bundle. In addition, the previous approaches do not account for potential mismatches between subjects, where features of one subject are not perfectly aligned with the features of other subjects. Furthermore, the models we fit in our previous work were linear models at their core, generalized only through the use of a link function that accommodated a limited family of non-linearities in the relations between brain tissue properties and phenotypical information.

The limitations of our previous approach are potentially overcome through the use of multi-layered neural network (NN) algorithms. This class of algorithms uses the backpropagation algorithm to fit a very large number of parameters, often organized into multiple layers with different operations happening within each layer and between layers. For example, a large sub-family of the broader class of algorithms are deep convolutional neural networks (CNNs), where the parameters of each unit within each of the convolutional layers of the model are restricted to perform a convolution operation over their inputs. These models, in particular, have been shown to be able to learn very non-linear relationships between their inputs and outputs, and to overcome significant systematic variance between training samples. For example, CNNs trained on classification of objects from photographic images can discriminate samoyed dogs from white wolves, even if they are photographed from different angles, or under a variety of illumination conditions ^6^. This suggests that CNNs would be able to take advantage of information that is spread over multiple locations in each bundle and over multiple bundles, as well as incorporate significant non-linearities into the analysis of tractometry data. On the other hand, because of the dependencies between neighboring nodes in a bundle profile in tractometry data, these data also resemble time-series data. Another family of algorithms, long short-term memory (LSTM) neural networks, which are a variant of recurrent neural networks (neural networks where information can be fed back into previous layers of the network), have been successfully applied to time-series analysis problems ^7^, suggesting that they might also offer some benefits in the analysis of tractometry data.

To test the performance of NN algorithms, compare different families of neural networks to each other, and compare them to a linear model, we conducted computational experiments using the Healthy Brain Network Processed Open Derivatives dataset (HBN-POD2; ^8^). This dataset contains dMRI measurements from more than 2,000 subjects, of which more than 1,800 have dMRI data of usable quality. As a benchmark of tractometry model performance, we used prediction of individual age. This task has three benefits as a benchmark: (i) age is a measurement with a ratio scale, meaningful units, and is estimated with very low or no error; (ii) brain white matter properties change during childhood and adolescence, providing a robust signal for prediction ^9,10^; and (iii) a wealth of previous research has demonstrated the value of prediction of individual age from brain data across a variety of settings ^11^. In addition to the direct comparison of model performance of our previous regularized regression approach, we also tested the data requirements of the neural network models, as well as the value of data augmentation, a method that injects noise into the samples used in training the neural network as a way of preventing overfitting to specifics of the training samples.

## Results

Model accuracy in age prediction was quantified using the coefficient of determination R^2^, which has a value of 0 for a model that does no better than predicting the mean age in the dataset for each subject, and 1.0, for a model that predicts accurately for every one of the subjects in the test set. To assess the impact of training set size, each NN algorithm and the baseline linear model were trained on a varying number of subjects. For each training set size, the models were each fit 10 times, allowing us to assess the reliability of model predictions. To facilitate comparison across models, the random seeds in each experiment were stored so that each model was tested on the same 10 different random samples of 20% of the subjects (n=264), and the training sets used in each of these 10 iterations of training were identical across the different models.

### Baseline model

Following our previous work^5^, we compared neural network performance to a baseline regularized linear regression algorithm: Principal Components Regression with Lasso regularization (PCR-Lasso). This algorithm starts by reducing the dimensionality of the data using principal components analysis (PCA) and then using the Lasso regularization approach to select the PCs that best fit the linear relationship with age. This is a strong baseline, as we have previously found this algorithm to reach state of the art performance on this task, with data very similar to the data used here. Indeed, this algorithm reaches maximal performance at R^2^ = 0.62 +/- 0.03 (SEM; Figure 1a). To characterize the dependence of algorithm performance on training set size, we fit an exponential learning curve model to these results (for example, in Figure 1a, the solid black line). The model fixes the maximal performance, based on the maximal performance observed in the data (α, Figure 3), and then fits two additional parameters: R^2^ at the smallest sample size (β, Figure 4) and the rate in which the curve between these two performance levels changes with increased training set size (*κ*, Figure 5). We find that PCR-Lasso improves substantially with the addition of training data. Performance in the smallest sample size (n=100) being R^2^ = 0.38 +/- 0.03 (SEM). Training set size dependence is such that ∼67% of the difference between the lowest and highest R^2^ (captured in *κ*) is achieved with 413 subjects.

**Figure 1:**
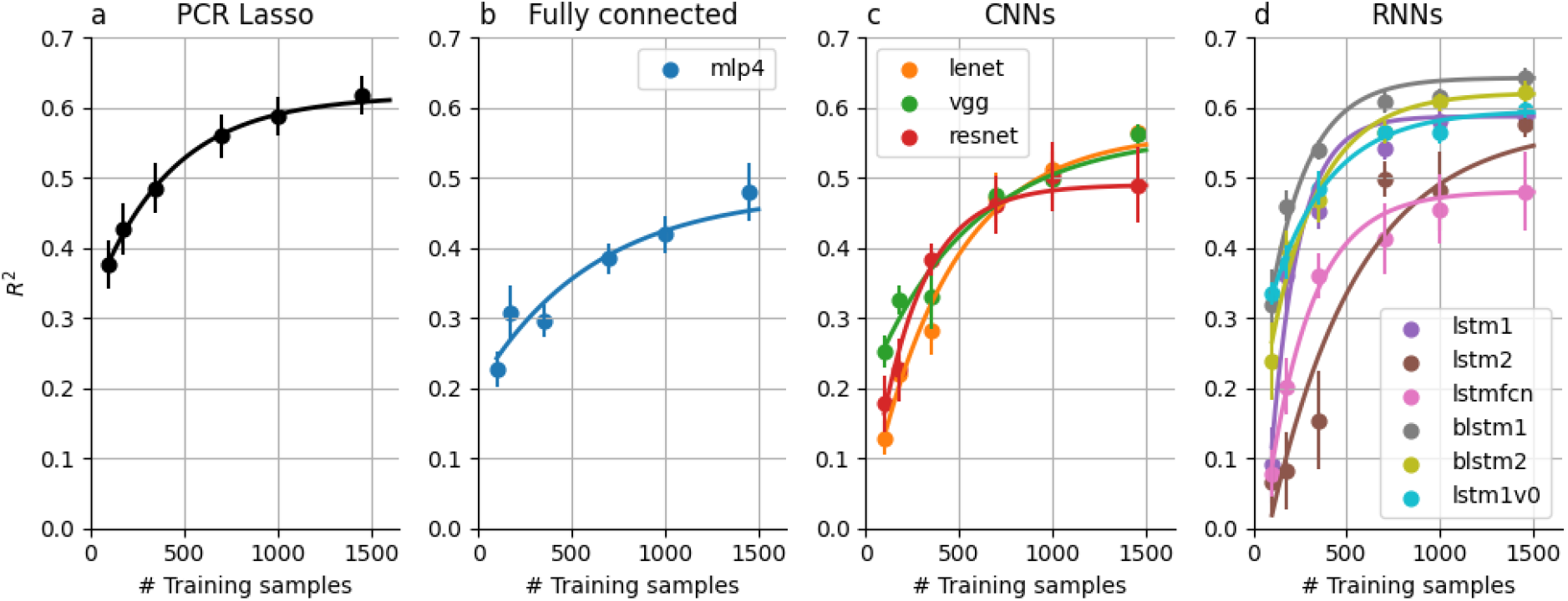
Performance of all brain age models increases with the number of subjects in the training set. (a) PCR Lasso, the linear baseline model starts at a relatively high R^2^, even at the smallest sample used here, and then increases from there. (b) MLP4, a fully connected algorithm has a similar learning curve, but it starts at much lower R^2^, (c) Convolutional neural networks (CNNs) start at an even lower R^2^ with small sample sizes, but increase precipitously reaching higher levels of performance at large sample sizes. (d) Recurrent Neural Networks (RNNs) have a variety of different performance characteristics, but are also overall very data hungry.

**Figure 2:**
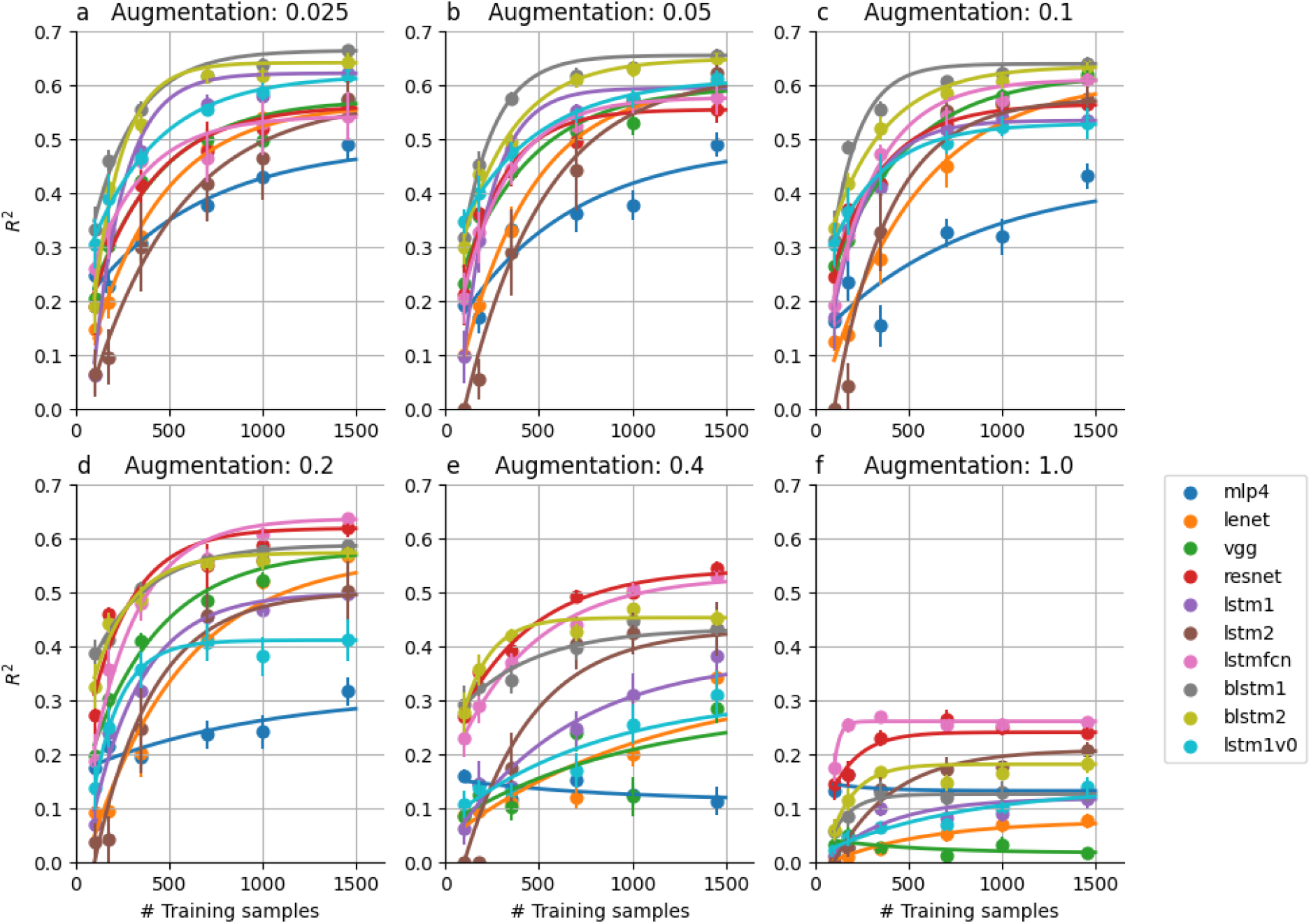
The effects of augmentation on NN algorithm performance. Several different levels of augmentation were applied (increasing from panel (a) through (f). While the performance of some models monotonically decreases with increased augmentation (e.g., mlp4, blue curves), some models’ performance increases with moderate levels of augmentation and then decreases when augmentation becomes too high (e.g., lstmfcn, pink curves). Further quantification of these trends is laid out in Figures 3, 4, 5.

**Figure 3:**
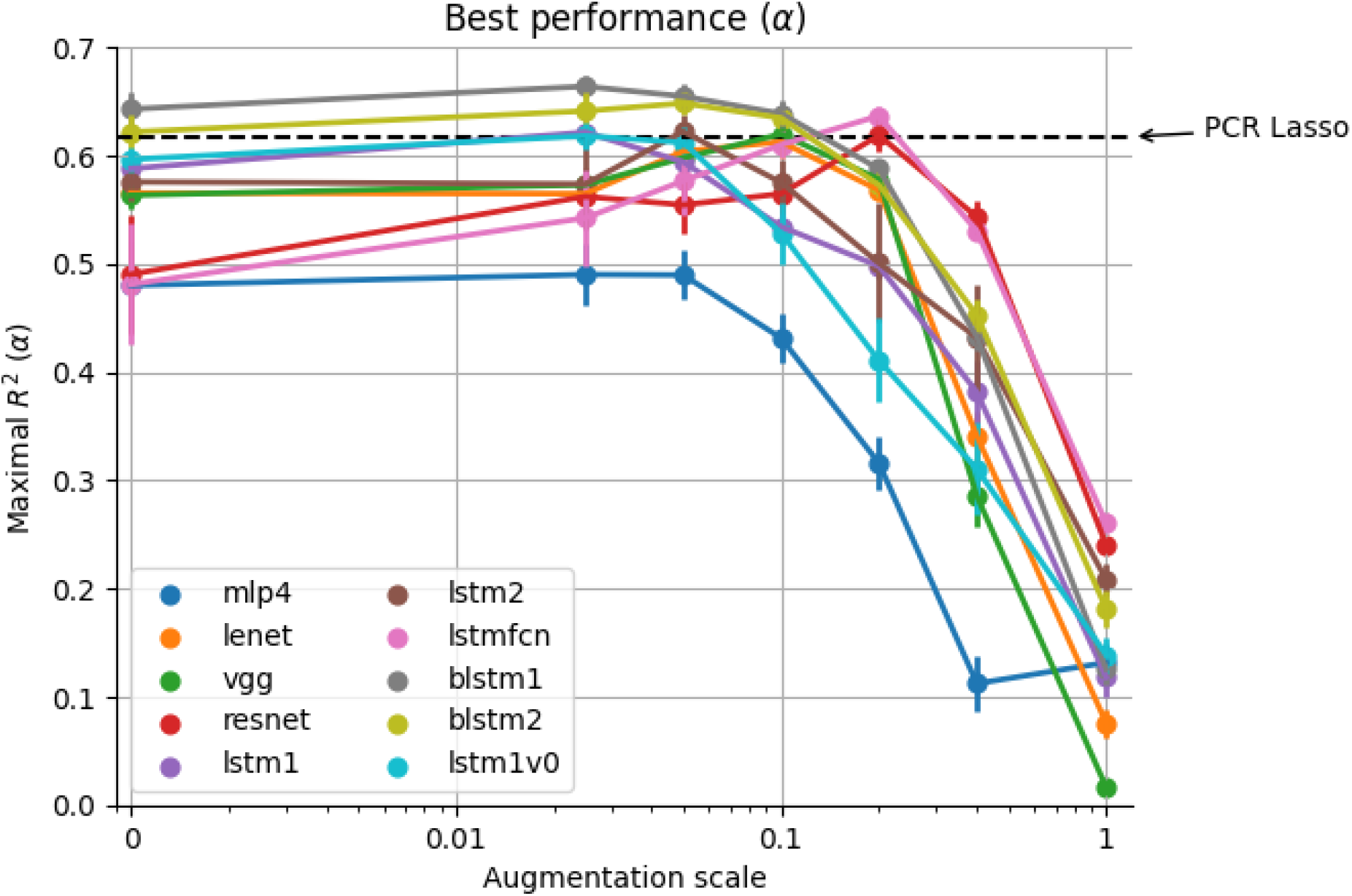
Best performance of the different algorithms. The black dashed line indicates the R^2^ of the linear baseline model (PCR Lasso). As seen in Figure 2, some algorithms only decrease in their performance with increased augmentation (e.g., mlp4, blue curve), but many of the NN algorithms improve their performance with increased augmentation, with some (e.g., resnet, red curve and lstmfcn, pink curve) reaching parity of R^2^ with PCR Lasso at higher levels of augmentation. At higher levels of augmentation the noise added to the measurements overwhelms all of the useful signal for training and all algorithms perform poorly.

**Figure 4:**
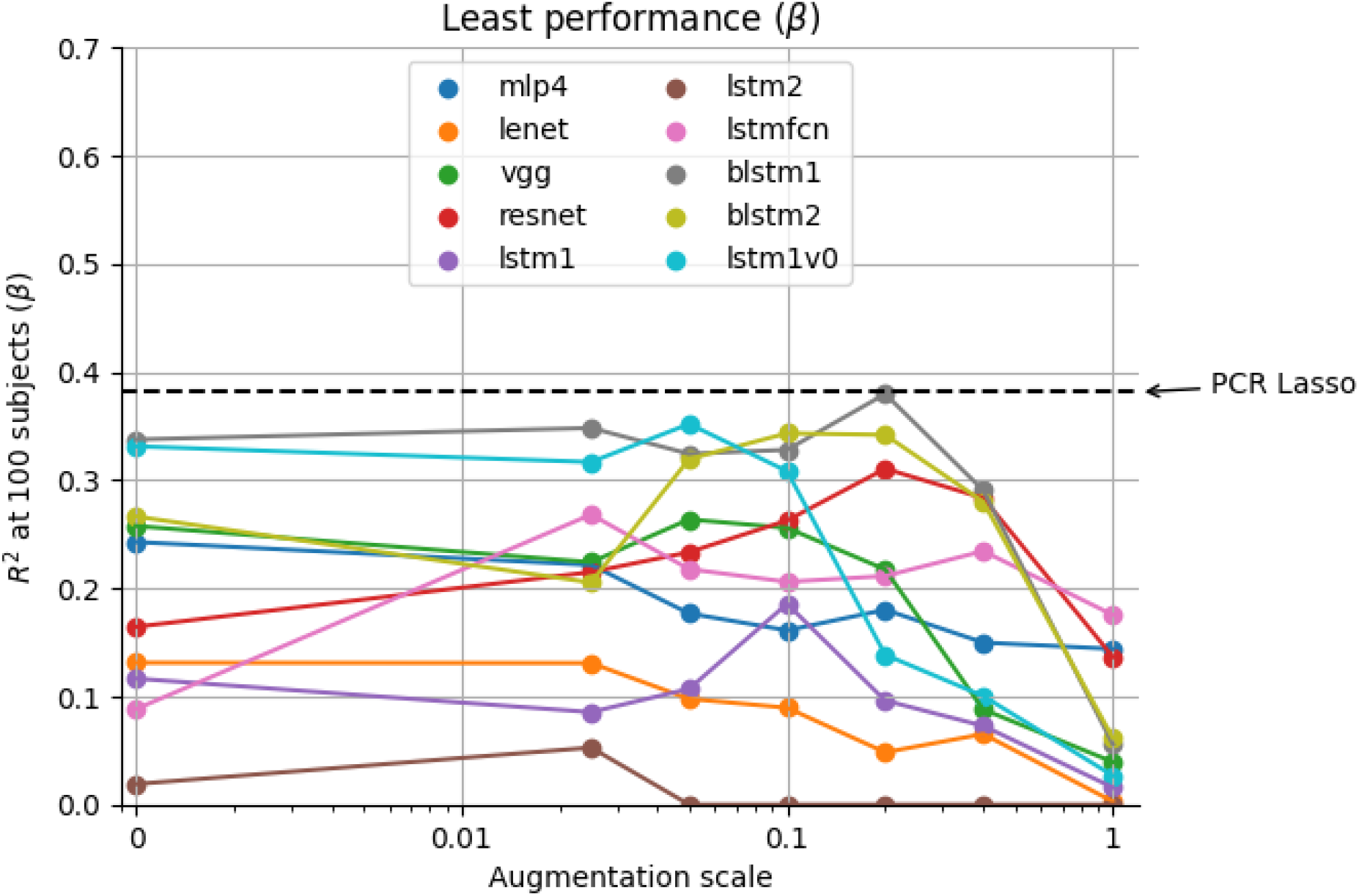
Least accurate performance. The resilience of the algorithms is quantified for cases where only limited data is available. Almost none of the algorithms, across all augmentation levels, are as resilient to smaller training data as PCR Lasso (dashed line).

**Figure 5:**
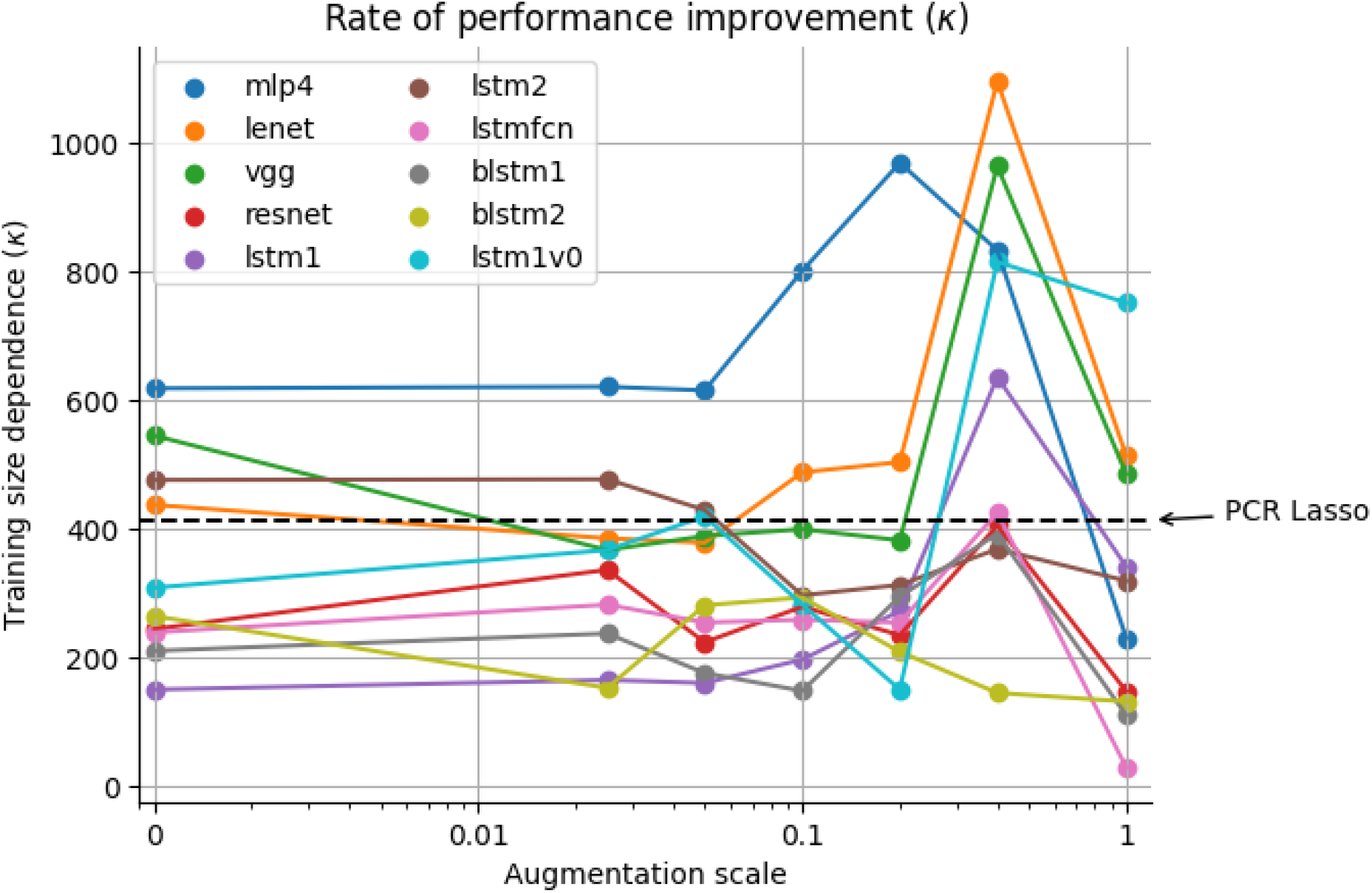
Rate of performance improvement with increased training sample size. Here, smaller values indicate more favorable performance (i.e., the algorithm requires a smaller sample size to reach ∼67% of its best performance). Several of the algorithms show improved resilience to small sample sizes, relative to the linear baseline (dashed line, PCR Lasso), even under conditions where performance is better than PCR Lasso (e.g., compare blstm1 curve with Figure 3).

### Neural network performance

As expected, NN algorithms are “data hungry”: all the algorithms exhibit poor performance at small sample sizes, with none of the NN algorithms reaching even the modest performance of the baseline model at this sample size (Figure 1). Nevertheless, we found that, for most of the NN algorithms, R^2^ reaches an asymptote within the range of training set sizes used here, but maximal performance varies widely. The mlp4, resnet, and lstmfcn algorithms all reach no better than R^2^=0.5, even when the largest training set is used (Figure 1b, c). On the other hand, some RNN algorithms reach parity with the baseline (blstm2; R^2^=0.62 +/- 0.016 SEM, Figure 1d), or even slight improvement above the baseline (blstm1; R^2^=0.64 +/- 0.015 SEM, Figure 1d). Importantly, some algorithms don’t seem to reach asymptotic performance, even with the largest sample size that we used, and their performance may improve with even larger samples. This seems particularly true for the more modestly sized CNNs: lenet and vgg (Figure 1c), as well as the lstm2 RNN algorithm (Figure 1d).

### The effects of data augmentation

Data augmentation introduces random noise to each sample of the training data in each batch of training. This method can help NN algorithms with a large number of parameters generalize better, by preventing the memorization of the samples in the training set. When augmentation levels grow very large, however, the signal in the data is overwhelmed by the noise that is added in augmentation, and the algorithm can no longer learn. In our data, we found that augmentation can have dramatic effects on algorithm performance in the brain age prediction task. For example, the resnet NN algorithm, which had poor R^2^ in the augmentation-free condition, reaches parity with the baseline model at relatively high augmentation levels (R^2^ = 0.62 +/- 0.016 SEM, Figure 3, red curve). The lstmfcn NN, which also performs poorly with no augmentation, reached even higher R^2^ than the baseline model with high levels of augmentation (R^2^=0.64 +/- 0.009 SEM, Figure 3, pink curve). However, at these higher levels of augmentation, the data requirements of these two models also increases (Figure 5). Algorithms that were similar in their performance to the baseline in the absence of augmentation improve slightly with the introduction of small amounts of augmentation. For example, the highest R^2^ reached by any model in these experiments is reached by the blstm1 model at a low value of augmentation (R^2^=0.66 +/- 0.01 SEM). The relatively-simple mlp4 model architecture that does not perform very well in the absence of augmentation, only becomes worse with the introduction of augmentation.

## Discussion

We found that brain age modeling can be more accurate with deep learning algorithms, but this advantage is small compared to regularized linear regression models. Our results suggest that DL approaches should be used with caution, and with comparison to strong linear baseline models, such as the PCR Lasso model that we used here. Moreover, as we posited in previous work ^5^, and consistent with previous findings ^9,10^, these results support the idea that brain development in childhood and adolescence is wide-spread throughout the brain and a process that occurs in tandem across large parts of the white matter. Under these conditions, non-linear and invariant modeling of the data may provide only minor benefits in terms of model accuracy. Nevertheless, some patterns do emerge from our analysis. In brain age modeling, RNNs and the blstm1 model, in particular, can emerge as superior to the linear baseline: this model has higher accuracy than the linear model at best, and its lowest R^2^ performance is not much worse than the linear baseline, while its rate constant of performance improvement is substantially smaller than the linear baseline, suggesting that it is even less data hungry than the linear baseline. This small improvement in R^2^ does come at the cost of substantially increased computational time, and reduced interpretability relative to the linear model (but methods for interpretation and explanation of complex non-linear models are also rapidly evolving to meet this challenge^12^). These considerations alone may steer researchers away from the use of deep learning algorithms in settings such as the one demonstrated here. In addition to these findings, CNNs, and particularly resnet and vgg, may gain in performance with increased sample sizes, as they do not seem to asymptote within the range of training sample sizes provided here, and they benefit from large amounts of augmentation.

What do these results tell us about the nature of tractometry data? On the one hand, as explained in the Introduction, the tractometry feature engineering pipeline produces tabular data, which should lend itself well to approaches such as PCR Lasso, which are not provided with a supervision signal about structural constraints of the data, such as spatial contiguity between different features, or their division into different groups of features (e.g., different white matter tracts). This is similar to many other machine learning problems where extracted features are not related to other features in any systematic way. On the other hand, tractometry data does have inherent structure: group structure, which we exploited in previous work^5^, as well as within-group spatial contiguity, which we exploit in the convolutional and recurrent NN algorithms used here. In fact, it is apparently not just the large number of parameters or capacity to represent non-linear relationships that provides a benefit to NN models: the one NN algorithm that performs most poorly is mlp4, which does have a large number of parameters, but is not designed to capitalize on the spatial contiguity of the data. More generally, when designing machine learning pipelines for analysis of large datasets in other domains, it will be important to take into consideration whether the algorithms that are used can take advantage of structure that is inherent to the data.

How general are these results for tract profile analysis of a range of different phenotypes? In other work, we found that deep learning methods may prove helpful in tasks that have different characteristics: In a study of a large group of participants with glaucoma and tightly-matched controls, we found that deep learning models (similar to the resnet model used here) can accurately classify glaucoma based on tractometry data from visual white matter pathways, while a linear baseline model (ridge-regularized logistic regression) cannot^13^. This means that the results presented here, while important in and of themselves, do not generalize to all machine learning tasks that researchers may choose to undertake with tractometry data, and to all samples. Linear models may work well with large signals (e.g., age), but it is possible that they are not as good at uncovering more subtle signals with complex non-linearities, as might be the case in many diseases. Hence, we hypothesize that NN models could have specific utility for uncovering subtle and complex signals associated with the early phases of disease, and conclude that NN modeling should be pursued with caution but also might be an important tool in the data-science toolkit for tractometry. Another future area of research where NN models that we used here may prove useful is in a generative setting. Previous work used a fully connected NN model, akin to the mlp4 model we used here, as the core of a generative normative model ^14^. Our results suggest that, provided enough training data, and with the benefits of data augmentation, normative modeling may benefit from some of the other architectures that we used here, such as CNNs and RNNs, a topic for further research.

In summary, our present results are important because: (i) brain age modeling is a commonly used ML task in neuroimaging; (ii) they lay out a framework for comparing different results across training data size and augmentation levels; and (iii) they provide an open-source software that other researchers can use in order to establish their own comparisons of different models applied to tractometry data in other studies. All of the neural network methods described here were implemented as part of an open-source software that we developed for multivariate tractometry analysis^5^, which is available at https://yeatmanlab.github.io/AFQ-Insight. More generally, our results serve as a warning to researchers across fields about the adoption of complex non-linear models in high-dimensional multivariate settings, where a high-performing regularized linear alternative would suffice. Future work across domains should focus on systematic analysis approaches – such as the one here – that defines the conditions under which different modeling approaches could be successful.

## Methods

### Data

We used data from the Healthy Brain Network ^15^ that we previously processed and automatically quality controlled ^16^. The measurements are described extensively elsewhere ^15,16^. Briefly, diffusion-weighted magnetic resonance imaging data was acquired with a spatial resolution of 1.8 × 1.8 × 1.8 mm^3^; 64 diffusion directions were measured with b=1,000 s/mm^2^ and b=2,000s/mm^2^ (in some rare cases, b=1,500 s/mm^2^ and b=3,000 s/mm^2^ were used). The data were curated using CuBIDS^17^ and then processed using QSIprep^18^. Quality control of the data was automated using a deep learning algorithm that was trained on community-scientist inputs, achieving high accuracy on a “gold standard” subset that was examined in detail by a group of experts ^8,19^. The original sample includes 2,747 subjects, aged 5-21 years, but after exclusion based on suitability for analysis and after selecting participants for which quality control was above 0, we included in the present study 1817 participants.

### Tractometry

We used automated methods to extract tract profiles of 24 major fiber bundles in every subject^20^. Each bundle was divided into 100 equidistant nodes, and dMRI-derived tissue properties were projected into each of these nodes. Tissue properties of the different bundles were characterized using the Diffusional Kurtosis Imaging model implemented in DIPY ^21,22^. To model the relationship between dMRI and age, we used fractional anisotropy (FA), mean diffusivity (MD) and mean kurtosis (MK) based on this model. Data were organized into a table in which each subject was a row, and each column represented one of these tissue properties in a node along the length of one of the 24 bundles.

### Baseline linear model

A baseline linear model was fit to the data. Based on our previous work on models of tractometry data ^5^, we used a PCR Lasso algorithm. In this algorithm the data dimensionality is first reduced using a principal components analysis, and then the Lasso regularization algorithm is used to select informative features ^23^.

### Neural networks

A variety of neural network architectures were implemented. We encoded the tract node as the “length” dimension in these one-dimensional networks and encoded the tracts/metrics as channels. Inspired by the use of neural networks in time-series classification, we borrowed heavily from the work of Iwana and Uchida^24^, who evaluated different data augmentation methods using ten different time series classification networks. These models can be composed into three broad groups which we describe below: fully connected networks, convolutional neural networks, and recurrent neural networks.

- Fully connected network: we use a fully connected multi-layer perceptron (MLP) model specifically tailored for time sequence labeling proposed by Wang et al.^25^. It ignores the structure of tract profile data by first flattening the input. It then passes the input through three hidden layers, each with 500 units. Dropout^26^ is added after the input layer, with a rate of 0.1, and after each hidden layer, with a rate of 0.2. We refer to this network as mlp4.
- Convolutional neural networks
  - 1-D Visual Geometry Group: Iwana and Uchida^24^ modified the original VGG^27^ network to accept one-dimensional time series input. They altered the number of convolutional blocks and max pooling blocks depending on the length of the input in order to prevent excessive pooling. We refer to this network as *vgg*.
  - LeNet-5: This 1-D CNN is an adaptation of LeCun et al.’s^28^ seminal network developed for handwritten character recognition. Similar to *vgg*, Iwana and Uchida^24^ altered the number of convolutional blocks depending on the length of the input to prevent excessive pooling. We refer to this network as *lenet*.
  - 1-D Residual Network: Here, we adopt the one-dimensional ResNet of Fawaz et al.^29^, who modified the original ResNet^30^ to contain only three residual blocks with varying filter lengths and no max pooling. We refer to this network as *resnet*.
- Recurrent neural networks
  - Long short-term memory (LSTM): We adopt three variants of the LSTM model. The first (*lstm1v0*) is an application of the original LSTM proposal^31^ consisting of a single LSTM layer with 512 units. The next two are adopted from a survey of LSTM hyperparameters for sequence labeling tasks^32^. These models have either one (*lstm1*) or two (*lstm2*) stacked LSTM layers, each with 100 units.
  - Bidirectional long short-term memory (BLSTM): these variants of the LSTM use both forward and backward recurrent connections. In the context of time series analysis, one can think of these connections as enabling prediction and retrodiction, respectively. We adopt two versions of the BLSTM, blstm1 and blstm2, which are identical to lstm1 and lstm2, except that they use bidirectional layers.
  - Long short-term memory fully convolutional network (LSTM-FCN): this model is a multi-stream neural network developed by Karim et al.^7^ combining an RNN and a CNN. The CNN stream consists of three 1-D convolutional layers with batch normalization, followed by global average pooling. The RNN stream consists of a single LSTM layer with 128 units and a dropout rate of 0.8. The outputs of these streams are then concatenated and fed into one fully connected node. We refer to this model as *lstmfcn*.

All of these models are implemented using the Keras API^33^ of TensorFlow 2^34^. Both the neural network code and the data augmentation code, discussed below, were inspired heavily by the implementations of Iwana and Uchida^24^, with some modifications for use on tractometry data and an adjustment to the output layer activation functions for use in regression.

### Data augmentation

We implemented a range of augmentation methods that are used in analysis of one-dimensional time-series^24^. In the experiments described here, we applied jitter, scaling, and time-warping. Each of these operations were parametrized by a single number which was scaled to the range of values within each bundle and metric (i.e, FA, MD and MK).

### Experimental design

Each model was evaluated for accuracy in age prediction 10 times. In each trial of the experiment, data were split into training (80%) and testing (20%) sets. The test set was set aside, to be used only in evaluating the trained network at the conclusion of training. We used the same test set, regardless of the size of the training data. Within each trial, different experiments were conducted with each of these test samples, where we sub-sampled the training data down to samples of size: 100, 175, 350, 700, 1,000 or the full training sample size of 1,453 subjects. Random seeds were fixed such that the same 10 test sets and the same training subsets were used with each model, facilitating the comparison across models. In cases where features could not be evaluated (e.g., because of localized issues with the dMRI data), data was imputed with a median imputation strategy applied within each set (train/test) separately.

For the baseline PCR Lasso model, the degree of regularization is set using an internal 3-fold cross-validation procedure within each training set.

For the NN models, Keras callbacks functions are utilized to stop training when there is no further improvement in performance and to store trained model weight (i.e., E`arlyStopping` with `monitor=`‘`val_loss`’`, min_delta=0.001, model=`‘`min`’`, patience=100; ReduceOnPlateau` with `monitor=`‘`val_loss`’`, factor=0.5, patience=20, verbose=1`). The ADAM optimizer ^35^ was used in all cases, and based on initial pilot experiments, we set slightly different learning rates for each NN architecture.

### Modeling training sample size effects

We fit the following exponential learning model to describe the effects of sample size on model performance:

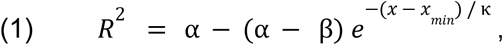

where *x* is the number of samples in the training set and *x*_*min*_ is the smallest training set used (here, *x*_*min*_ = 100 in all cases), α is the highest R^2^ observed in the data and is fixed *a priori*. Two parameters are fit: β is an estimate of the performance that would be achieved at the smallest training sample, *κ* is a parameter that quantifies the rate at which performance improves with additional data. Given the simple exponential form of this curve, this parameter corresponds to the number of subjects at which performance improves to 67% of the difference between β and α. The model was fit to the data using the Scipy software library’s ^36^ `optimize.curve_fit` function.

Code running all of these experiments, modeling, and visualizing their results is available in https://github.com/rlqiao/afq-deep-learning/. The neural network implementations and augmentation methods are implemented as part of the open-source software toolbox “AFQ-Insight” available at https://yeatmanlab.github.io/AFQ-Insight.

## Acknowledgements

This research was supported through grant RF1MH121868 from the National Institutes for Mental Health/The BRAIN Initiative. Cloud computing was provided via the University of Washington eScience Institute Azure Cloud Compute Credits program and through the Amazon Web Services Cloud Credits for Research program. Thanks go to the University of Washington eScience Institute core staff for helpful discussion of these results.

## Author contributions

All authors conceived the experimental design. ARH implemented neural network and augmentation software. AR, ARH, and JQ conducted computational experiments. AR analyzed experimental results. AR wrote the initial version of the manuscript. All authors revised the manuscript. AR and JDY acquired funding.

## Data availability statement

All of the MRI data used here is openly available to use at https://fcp-indi.s3.amazonaws.com/index.html#data/Projects/HBN/BIDS_curated/, and fully described in ^8^. Code to run the experiments described here and to visualize the results is available in https://github.com/rlqiao/afq-deep-learning/.

